# Spatial heterogeneity and evolutionary dynamics modulate time to recurrence in continuous and adaptive cancer therapies

**DOI:** 10.1101/128959

**Authors:** Jill A. Gallaher, Pedro M. Enriquez-Navas, Kimberly A. Luddy, Robert A. Gatenby, Alexander R. A. Anderson

**Affiliations:** H. Lee Moffitt Cancer Center, Integrated Mathematical Oncology (Tampa, FL, United States); H. Lee Moffitt Cancer Center, Cancer Imaging and Metabolism (Tampa, FL, United States)

## Abstract

Treatment of advanced cancers has benefited from new agents that supplement or bypass conventional therapies. However, even effective therapies fail as cancer cells deploy a wide range of resistance strategies. We propose that evolutionary dynamics ultimately determine survival and proliferation of resistant cells, therefore evolutionary strategies should be used with conventional therapies to delay or prevent resistance. Using an agent-based framework to model spatial competition among sensitive and resistant populations, we apply anti-proliferative drug treatments to varying ratios of sensitive and resistant cells. We compare a continuous maximum tolerated dose schedule with an adaptive schedule aimed at tumor control through competition between sensitive and resistant cells. We find that continuous treatment cures mostly sensitive tumors, but with any resistant cells, recurrence is inevitable. We identify two adaptive strategies that control heterogeneous tumors: dose modulation controls most tumors with less drug, while a more vacation-oriented schedule can control more invasive tumors.

## Introduction

Despite major advances in cancer therapies, most metastatic cancers remain fatal because tumor cells have a remarkable capacity to evolve drug resistance, both through genetic and non-genetic mechanisms (1). Most investigations of cancer treatment resistance have focused on identifying and targeting the molecular mechanisms that confer resistance. However, defeat of one resistance strategy often results in the deployment of another (2).

An alternative approach focuses on the population-level dynamics governed by Darwinian evolutionary principles that define the fitness of each cell within the local environmental context. For example, cancer cells often employ multidrug resistance pumps, in which the synthesis, maintenance, and operation require considerable investment of resources (up to 50% of the cell’s total energy budget) (3). In the harsh tumor microenvironment this investment in survival will likely require diversion of resources that would ordinarily be devoted to invasion or proliferation. Thus, while tumor cells may possess the molecular mechanisms necessary for therapy resistance, proliferation of resistant cells is governed by complex interactions that include the cost/benefit ratio of the resistance mechanism(s) and competition with other tumor subpopulations.

A common maxim in cancer treatment is to "hit hard and fast" through maximum dose dense strategies that administer the highest possible drug dose in the shortest possible time period. The maximum tolerated dose (MTD) principle has been the standard of care for cancer treatment for several decades and is the basis for clinical evaluation for most Phase I cancer drug trials. It has not, however, resulted in consistent cures in patients with most disseminated cancers (4). An evolutionary flaw in this strategy is the assumption that resistant populations are not present prior to therapy. It is now clear that cancer cells can be insensitive even to treatments that they have never “seen” before. Therefore, MTD therapy designed to kill as many cancer cells as possible, although intuitively appealing, may be evolutionarily unwise. This is because of a well-recognized Darwinian dynamic from ecology termed “competitive release”. Observed, for example, when high doses of pesticide are applied for pest eradication (5), competitive release allows rapid emergence of resistant populations because of the combination of intense selection pressure and elimination of all potential competitors.

Despite the growing recognition that heterogeneity and evolution play a significant role in driving treatment failure, explicit inclusion of Darwinian principles in clinical trial design is rare (6–9). However, both clinical and pre-clinical studies have shown promising results. Enriquez-Navas et al. used an evolution-guided treatment strategy to control breast cancer tumors in mice (10). They found that progression free survival can be prolonged when paclitaxel treatment schedules incorporate dose modulations and treatment holidays such that less drug is given to a responding tumor and more to a rebounding tumor (Fig. 1A). An ongoing clinical trial at Moffitt Cancer Center (NCT02415621) tests these evolutionary principles in patients with metastatic castration resistant prostate cancer to try to prevent the evolution of resistance to abiraterone therapy (11). In this trial, abiraterone is discontinued when the blood Prostate Specific Antigen (PSA) concentration falls below 50% of the initial value and does not resume until the PSA returns to the pre-treatment level (Fig. 1B). It is important to note that: 1) each patient serves as their own control to calibrate the PSA as a relative value, and 2) the adaptive schedules effectively personalize the treatment to patient response so that while one patient has only 2 courses of treatment in a year, another gets 3 in 10 months. Yet both remain under control.

**Figure 1.**
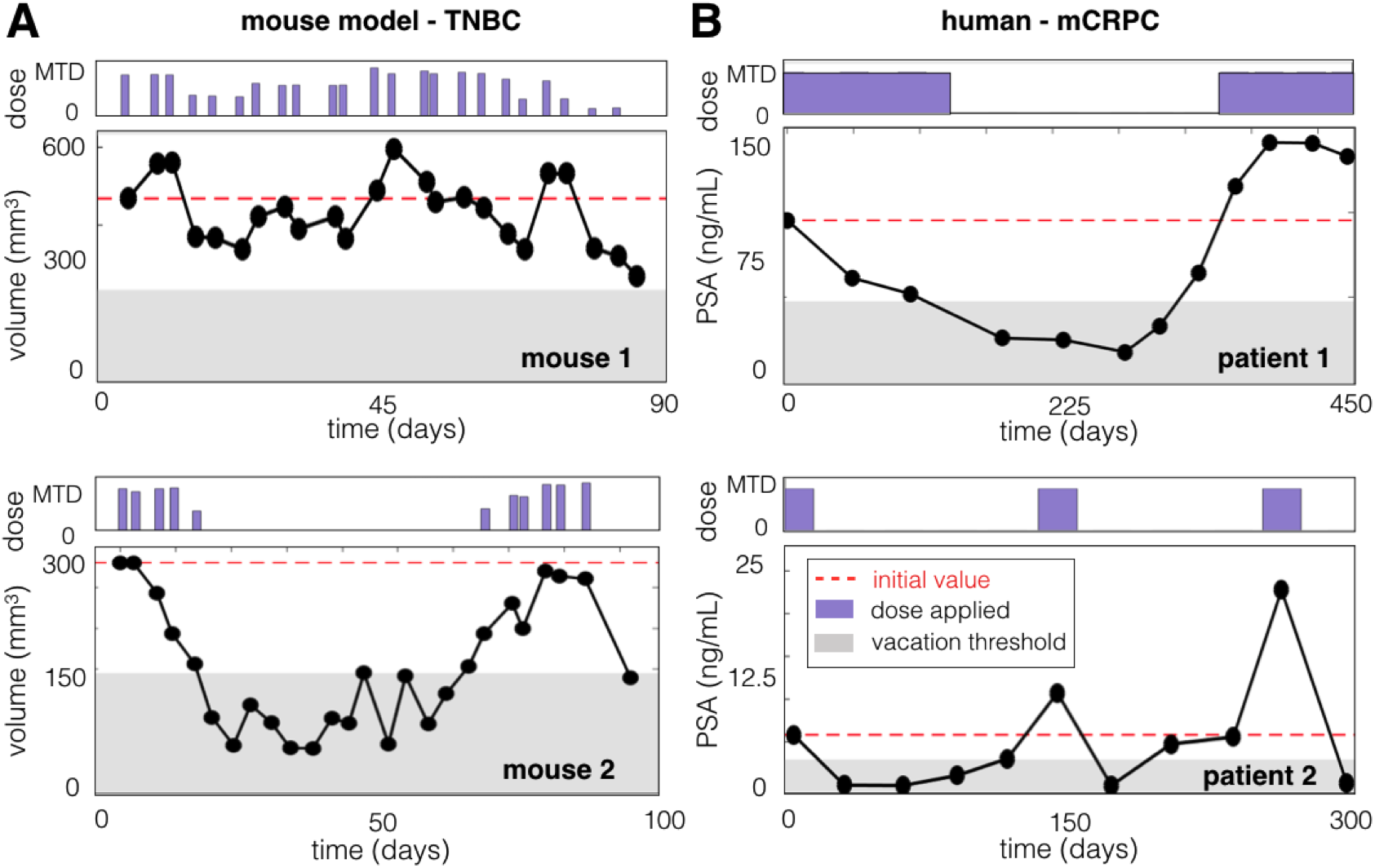
Adaptive therapy in the laboratory (A) and the clinic (B). A) Mice implanted with triple negative human breast cancer were treated with adaptive paclitaxel treatment. If the volume dips below 150 mm^3^, a treatment vacation occurs, if there is a 20% tumor volume decrease (or increase), there is a 50% dose decrease (or increase), and otherwise the dose remains the same (see (10) for details). B) Patients with metastatic castrate resistant prostate cancer are treated with abiraterone such that treatment is stopped if PSA falls below 50% of the original and resumes when the PSA exceeds the original value.

Two important questions emerge from these results: (i) For which cancers is continuous (MTD) treatment the best strategy, and when is adaptive better? (ii) For adaptive treatments, when should a patient receive treatment holidays instead of dose modulation? While selection for resistance through application of continuous cytotoxic therapy seems inevitable, proliferation of those cells may be controlled using evolutionary principles. Importantly, multiple experimental models have shown that drug-resistant cancer cells proliferate slower than sensitive cells in the absence of drug (12–16). This is because resistant tumor cells, like most drug-resistant bacteria, incur a fitness cost due to the energy costs involved (17). Here, we computationally investigate these questions under the hypothesis that these costs can be exploited to delay or prevent proliferation of resistant cells in the tumor.

## Materials & Methods

To investigate the effect of spatial competition and heterogeneity in tumor populations under different drug schedules, we modify a 2-dimensional off-lattice agent-based model (described previously in (18)) to incorporate treatment response. The model flowchart is shown in Fig. S1.

### Off-lattice agent-based model

We use an off-lattice agent-based model to investigate the evolutionary dynamics of different treatment schedules on heterogeneous tumors. Space is a limiting factor, such that at carrying capacity, a cell will enter quiescence due to contact inhibition. Space is deemed available if any contiguous set of integer angles are empty and sufficient to allow a cell to fit without overlap (described previously in (18)). When a cell is quiescent, we assume that it cannot proliferate and is not affected by the drug. We also assume that there is no cell death due to regular turnover, and resources are abundant when cell densities are below carrying capacity. Model flowchart is shown in Fig. S1.

### Tumor growth model

We begin with a small mass of 100 cells clustered in the center of the domain. The simulation is stopped when the number of cells reaches 15,000, which produces a mass around 1.5 mm in diameter, representing a micrometastasis.

### Treatment

For continuous therapy, we apply the maximum tolerated dose (MTD) the entire duration, but only when a cell is capable of dividing does it become sensitive to drug toxicity. For adaptive therapy, each dose is instantaneously realized, ignoring any short-term pharmacokinetics. Regardless of the treatment schedule, a cell’s response to drug exposure is defined as a probability of death, *P*_*death*_, which depends on its sensitivity, *s*, and the dose, *D*:

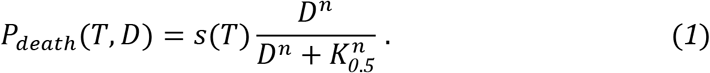

Due to the resistance cost, we assign the cells with the shortest cycle times, *T*_*min*_, 100% sensitivity, and in a linear fashion, the longest cycle times, *T*_*max*_, are completely resistant, i.e. *s(T)*=*1-(T-T_min_)*/*(T_max_-T_min_)*. This provides a simple linear trade-off between a cell’s fitness in the absence and presence of drug (12–16). We use a Hill function to describe the dose effect (19,20), setting the Hill coefficient and half-maximal activity to *n*=1.5 and *K*_*0.5*_=0.125. These values are not specific, but generically assume a response function that gives nearly 100% probability of death at the highest drug concentrations, dropping off slowly at mid ranges and then quicker as it gets to lower concentrations. The dose varies from its previous value *D*_*0*_ based on the relationship of the current number of cells *N* to the previous number of cells *N*_*0*_ according to the following equation:

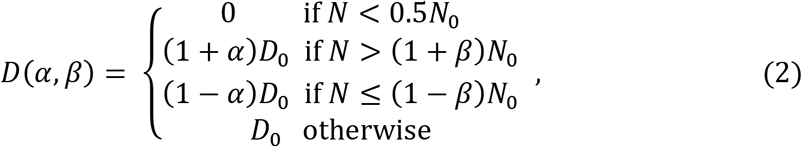

where *α* and *β* are fixed values that affect how much the dose changes due to population size changes. In Eq. 2, if the number of cells is below half of the original, a treatment vacation occurs. If there is a fractional increase in number of cells greater than *β*, there will be a fractional increase in the dose by *α*, and if there is a fractional decrease in population size greater than *β* there is a fractional decrease in dose by *α*. Otherwise, the dose remains the same. The dose always begins at the MTD, never exceeds the MTD, and each new dose is determined every 3 days in accordance with the experiments reported in (10). If a cell is targeted by the drug, there is a 15-30h delay, randomly chosen, before it is removed from the domain, to account for the time it takes for apoptosis to occur (21,22).

Treatment is stopped when either the tumor is cured (i.e. the number of current cells *N*=*0*), recurs (i.e. *N*=*4/3N_0_*=20,000 cells), or the tumor reaches an age of 2 years post-treatment. We assume tumor control if the final number of cells is below 10,000, which is half the value that determines recurrence and accounts for fluctuations during adaptive treatments.

### Cell migration

To examine cell migration, we assume all cells move in a persistent random walk. Each cell is given a persistence time (randomly chosen from a normal distribution with 80±40 minutes) during which it will move. After this time, it will turn at a random angle and start again. All cells move as long as they are not in the quiescent state and don't contact another cell. Upon contact, the cells turn by a random angle and start again with a new persistence time.

### Phenotypic drift

We examine phenotypic drift by allowing daughter cells to inherit a slightly different proliferation rate upon division. At each division, there is a 10% probability that a cell will randomly choose 1 of 3 options: increase its cycle time by 1 h, decrease its cycle time by 1 h, or stay the same. The cell cycle is bounded within the range of 10-50h.

## Results

### Fitness differences and space limitations affect competition and selection

The cost of resistance and the selection force imposed by space limitations can be demonstrated through a combination of *in vitro* and *in silico* models. The MCF7Dox cell line is highly resistant to many chemotherapy agents due to upregulation of the membrane efflux pump P-glycoprotein or Multiple Drug Resistance (MDR1) proteins. These cells were labeled with red fluorescence protein (RFP) and initially plated at a 1:1 ratio with the parental MCF7 cell line labeled with green fluorescent protein (GFP), which does not express MDR1 and remains sensitive to chemotherapeutic agents. In these experiments, the cells are grown for 3-4 days, harvested, counted, split and re-plated over 46 days. Each cell type was measured by flow cytometry at each interval (top left Fig. 2). The resistant cells, because of the fitness cost of the MDR expression, are rapidly outcompeted by the parental MCF7 cells after only a few generations in the absence of chemotherapy (doxorubicin) in the media.

**Figure 2.**
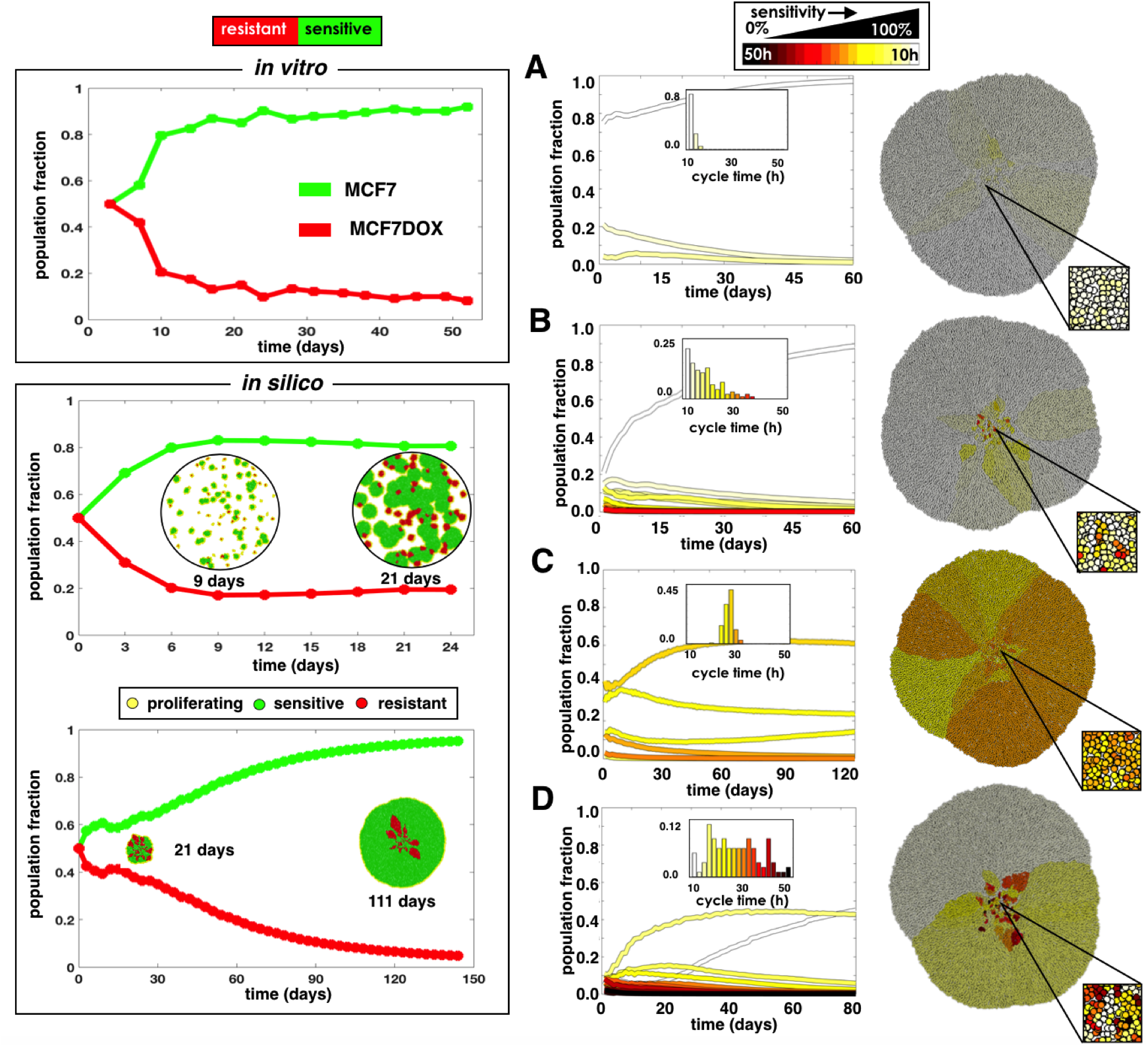
Competition between sensitive and resistant cells when populations are binary (left) and heterogeneous (right) using *in vitro* and *in silico* models. Cells sensitive (MCF7) and resistant (MCF7Dox) to doxorubicin are co-cultured *in vitro* and replated every 3-4 days (top left). The plot shows over time the fraction of drug-sensitive cells outcompetes drug-resistant cells. An *in silico* co-culture of sensitive (40h cell cycle) and resistant (60h cell cycle) cells seeded randomly throughout the domain or clustered in the center (lower left). Both simulations were initialized with 100 cells at a ratio of 1:1 and grown to 15,000 cells. The spatial distribution of cells is shown at several time points. A-D) *In silico* simulations were initialized with different phenotypic distributions, characterized by a mean sensitivity s and standard deviation σ_s_: A) s=100%, σ_s_=5%; B) s=100%, σ_s_=25%; C) s=60%, σ_s_=5%; and D) s=60%, σ_s_ =25%. These initial distributions are show as histograms in the plot insert; the final spatial layout is shown to the right.

We then investigated these evolutionary dynamics through our computational model system. We assumed that the population mix consists of a faster growing sensitive phenotype and a slower growing resistant phenotype. We use the average cell cycle times for the sensitive and resistant cells found at the beginning of the *in vitro* experiment (MCF7: 40h, MCF7Dox: 60h). We initially seeded 100 cells in 2 configurations: randomly scattered throughout a 1.5mm radius circular domain, representing an *in vitro* cell culture distribution, and tightly clustered in the center, representing a dense tumor mass (lower left Fig. 2). With the scattered distribution, due to the low density, there is ample space for most cells to grow unimpeded. While there is selection for faster growing sensitive cells, each colony has its own “edge” where cells remain proliferating. In contrast, when cells are clustered closely together, they are less free to move and proliferate. Proliferation is thus confined to a narrow advancing edge, so the population grows much slower. This adds additional selection pressure, which causes sensitive cells to quickly take over the expanding front and trap the resistant cells within.

Sensitivity to drugs is often viewed as binary, where a cell that is exposed to drug simply dies if sensitive or survives if resistant. However, in reality tumors have a more nuanced mix of phenotypes. We investigate how various mixtures of sensitive and resistant cells compete to form a solid tumor mass, starting with 100 cells with cell cycle times randomly drawn from a normal distribution characterized by a mean, *s*, and a standard deviation, *σ*_*s*_. Figure 2A-D shows the final spatial configuration of some tumors grown from different initial populations. In each case, the more resistant cells get trapped in the interior of the tumor by the more sensitive cells, which take over the invading edge. Coexistence between different phenotypes is seen, but given enough time the more proliferative phenotypes take over the invading edge. Overall, the trend shows that for a heterogeneous distribution of phenotypes, the faster proliferators will dominate the population, but the timing will depend on the relative fitness differences.

### Continuous treatments can cure some tumors; adaptive treatments can control most tumors

We compare a conventional continuous MTD treatment strategy (CT) with an “adaptive therapy” (AT), wherein we adapt the next treatment based upon the tumor’s previous response. We use an adaptive scheme that will change the dose by 25% (*α*=0.25) if the population size changes by 5% (*β*=0.05), applies a treatment vacation if the population is below half the original, and otherwise stays the same (see Eq. 2). Treatment is applied to the tumors from Fig. 2A-D until either the tumor is cured, recurs, or the tumor reaches an age of 2 years posttreatment.

In general, we find that there is not one therapy strategy that works best for all tumors, but the response to the treatment depends on the tumor composition (Fig. 3). For the most sensitive and homogeneous tumor (Fig. 3A), CT can kill all cells completely while AT can keep the tumor under control by using an average dose of 34% of the MTD. While the total dose for AT is ~2 times that of CT, it is distributed over a time period ~6 times as long. The tumor consisting primarily of sensitive cells, but includes some treatment-resistant phenotypes (Fig. 3B), initially responds very well to CT, but this is followed by recurrence of a completely resistant population after ~1 year. The same tumor under the AT can be controlled with an average of only 35% of the MTD with a total dose that is 70% of that given using CT. The more resistant tumor (Fig. 3C) should yield a slight net negative growth rate with the MTD applied. However, after a modest initial response to CT treatment, resistant cells take over enough to yield a positive net growth rate, which leads to tumor progression. Using the AT strategy, the change in the population is too slow to induce dose modulation, however, when the population halves at around 1 year, a treatment vacation is applied. At this point, most of the sensitive cells have already been eliminated and the tumor is spatially diffuse. Consequently, a large portion of actively proliferating cells quickly expand, and the tumor recurs 1 month prior than when using the CT strategy. For the more resistant and heterogeneous tumor (Fig. 3D), both treatments lead to eventual recurrence, but the total number of cells is kept stable for a while using AT. Notably, AT gives ~2 times the dose but over a time period that is ~2.4 times longer. Recurrence is delayed using an average of 79% of the MTD.

**Figure 3.**
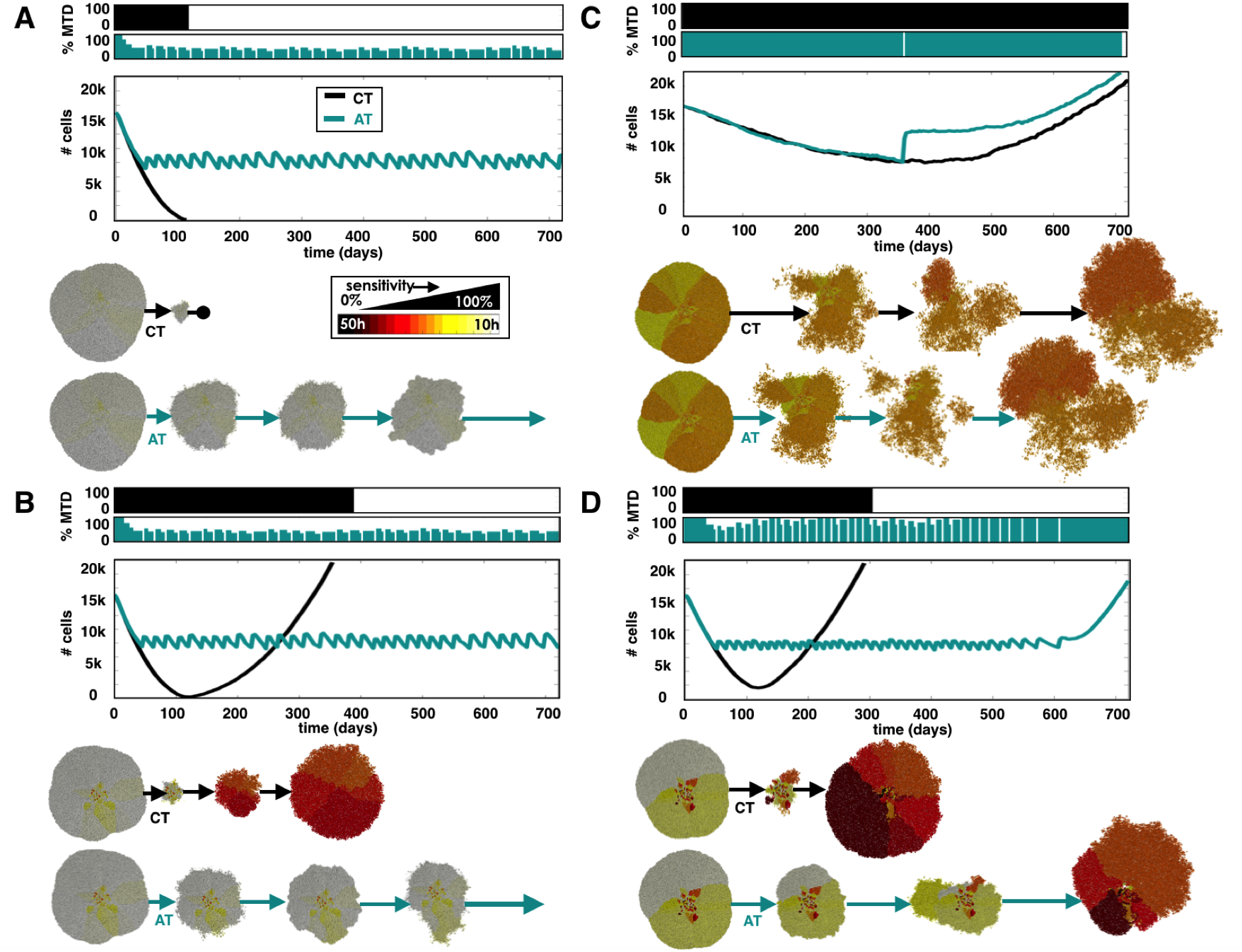
CT and AT schedules applied to different compositions of cell phenotypes, with mean sensitivity s and standard deviation *σ*_s_: A) *s*=100% and *σ*_*s*_=5%, B) *s*=100% and *σ*_*s*_=25%, C) *s*=60% and *σ*_*s*_=5%, and D) *s*=60% and *σ*_*s*_=25%. Top panels show the dose schedules for each treatment strategy, middle panels, the population dynamics, and lower panels, the spatial configurations at several time points. See link in the Supplement for animations of each of these different treatment strategies.

### A more vacation-oriented strategy can control the tumor at the expense of higher doses

The AT schedule used so far, which we now refer to as AT1 (*α*=0.25, *β*=0.05), was chosen somewhat arbitrarily. Changing *α* and *β* changes the amount the dose is modulated (*α*) if the population changes by a threshold amount (*β*), therefore these values can have a direct impact on AT control. We test another schedule that boosts both values, requiring a larger change in population size (10%) to produce a larger change in the dose modulation (50%), which we will call AT2 (*α*=0.50, *β*=0.10).

We compare all treatment strategies (CT, AT1, and AT2) in Fig. 4 using a tumor with a pre-growth distribution that centers around a sensitive phenotype with a large degree of heterogeneity (*s*=100%, *σ*_*s*_=25%). At the start of treatment the resistant cells are trapped in the interior by sensitive cells. The tumor recurs with the CT strategy after a year while both AT strategies control the tumor for the full 2 years (Fig. 4A). We find that the AT1 dose lowers at the start then incorporates both vacations and dose adjustments to control the tumor using an average of 35% of the MTD. In the AT2 schedule, however, we find that the population does not change sufficiently fast to invoke a dose change, so control is achieved solely by having treatment vacations giving an average of 76% of the MTD. However, since the proliferating cells are very sensitive to the drug, a large dose is not needed to control the tumor, so the more modulating AT1 dose schedule gives less drug to the patient while achieving the same effect. The average sensitivity vs. the average dose for each schedule is plotted in Fig. 4B against a background heat map of the expected net proliferation rate of the tumor 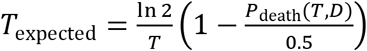. With CT, the mean sensitivity decreases over time as the most sensitive cells are eliminated and the more resistant cells take over. AT schedules, however, keep the cells, on average, in the sensitive region. The AT1 strategy shifts to the lowest dose that still kills the most sensitive phenotypes while the AT2 dose remains high.

**Figure 4.**
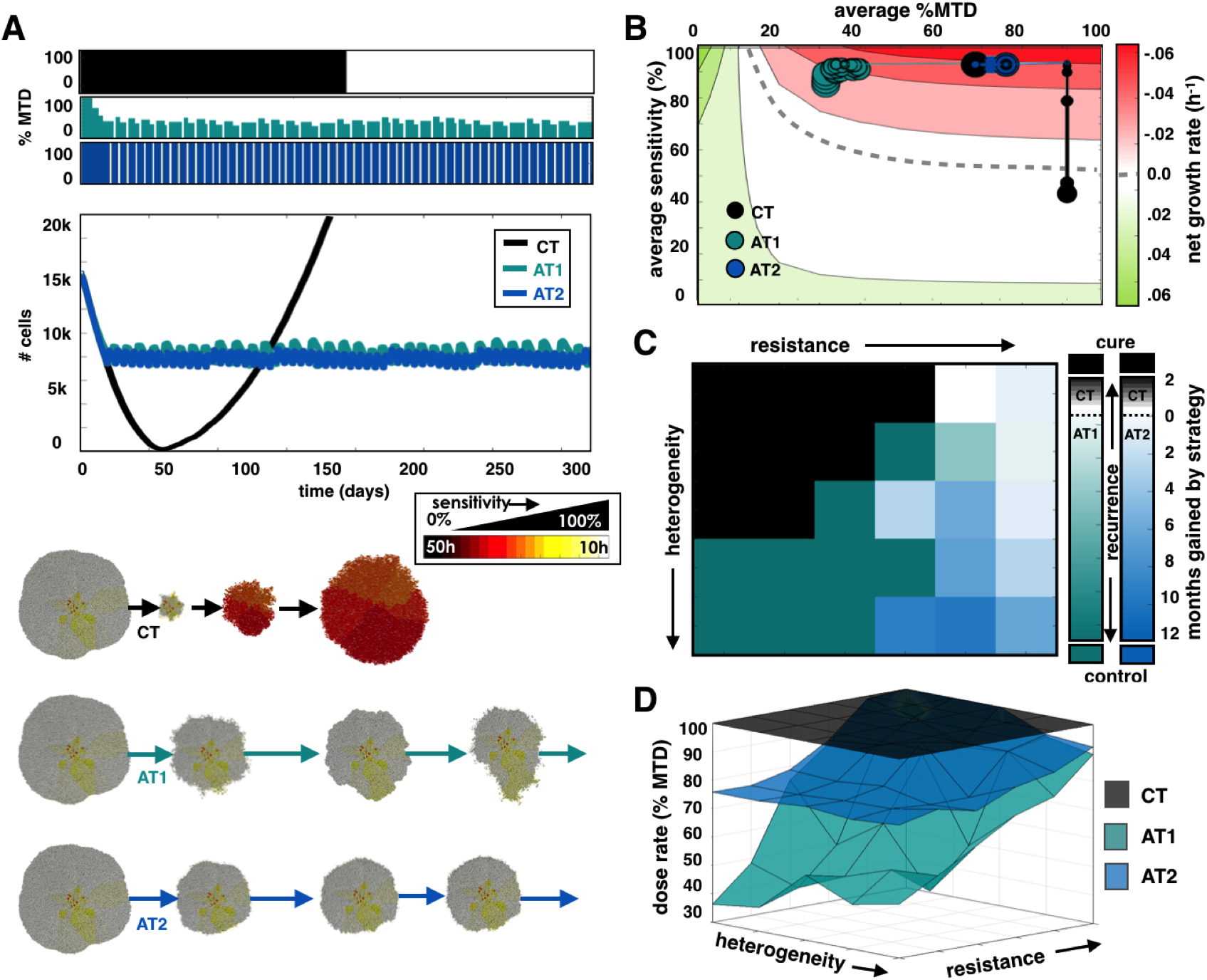
A) Comparing CT, AT1 (*α*=0.25, *β*=0.05), and AT2 (*α*=0.50, *β*=0.10) treatments using a tumor from a pre-growth normal distribution of *s*=100% and *σ*_*s*_=25%. Dose schedules, population dynamics, and spatial layout at various time points are shown for each treatment. See link in the Supplement for animations of each of these different treatment strategies. B) Trajectories for average sensitivity vs. average dose plotted every month with increasing point size. The background color indicates the expected growth rate of a tumor with a given cell cycle time, T, receiving a drug dose D (see text). The dashed line indicates where net growth is expected to be zero. C) Heat map indicates the strategy that gives the best outcome averaged from 3 simulations for different tumor compositions. Outcomes are favored in following order: cure, control, and longest time to recurrence. If recurrence is most likely, color indicates the average time gained by the winning strategy. D) Average dose given for each treatment strategy.

We can compare these 3 strategies over a range of different tumor compositions to determine which is most efficacious in each case. Specifically, we grow and test treatments on an array of tumors with different starting distributions with unique sensitivity and standard deviation (*s*=50-100%, *σ*_*s*_=5-25%). Presuming that the descending order of desired outcomes is to: cure, maintain, then gain the most time before recurrence, we obtain the best choice of treatment using 3 trials for each tumor composition in the array (Fig. 4C). When several strategies give the same results, preference goes to the lowest dose. We find that the most sensitive and homogeneous tumors are cured by CT, but AT strategies do better for heterogeneous tumors. The AT1 schedule provides a lower dose when the cells are more sensitive and controllable, the AT2 schedule extends the time to recurrence when recurrence is inevitable, and the more resistant, less heterogeneous tumors recur similarly regardless of treatment strategy. The time to recurrence using each treatment schedule over the array of tumors is plotted in Fig. S4A. The average dose given for the array of tumor compositions are given in Fig. 4D for each treatment. We find that we can control tumors bounded by drug-sensitive cells either by shifting to lower doses (AT1) or applying vacations between high doses (AT2). They both work because very sensitive proliferating cells do not need a full dose to be killed. However, when the tumor contains more resistant phenotypes, not all cells will respond to lower doses, so high doses and vacations grant better control.

### More treatment vacations are needed to control heterogeneous invasive tumors

For the adaptive schedules to work, the sensitive cells must impede the proliferation of the resistant cells by competing for space and trapping the resistant cells inside the tumor. However, if the cells can move, the spatial structure, that keeps the cells quiescent and hidden from the drug, is disturbed. We now examine the effect of cell migration by allowing cells to move in a persistent random walk at a modest speed of 5 μm/h (see methods for details).

Figure 5A shows an example with the same initial conditions as the previous example (*s*=100% and *σ*_*s*_=25%), but with migrating cells. The tumor composition appears similar to the previous example, but the cells are more spatially mixed. In this case, both the CT and AT1 schedules lead to recurrence while the AT2 schedule maintains control for the full 2 years (not fully shown). We see that for the CT case, compared to the case without migration (Fig. 4A), we have quicker decline in the population followed by faster relapse at 153 days. Due to the spatial spreading from migration, less cells are quiescent so the drug is able to affect more cells. This also allows resistant cells to escape confinement sooner, leading to a faster recurrence. For AT1, dose modulation occurs, but the tumor population doesn't get small enough to trigger a vacation. The dose reduces substantially, and then rises again when resistant, unresponsive cells take over. The AT2 dose schedule incorporates treatment vacations, but also modulates because the larger proliferating fraction of cells (that are susceptible to the drug) causes larger fluctuations in the population size during both growth and treatment phases, triggering dose changes. Importantly, by employing the quicker switching between fast growth (during the vacations) and fast death (many cells susceptible and higher doses), AT2 keeps the sensitive population at the invasive edge and the resistant cells suppressed. The AT2 schedule gives a larger average dose (67% of the MTD compared to 30% of the MTD for AT1), but does not recur over the monitored 2 year time period. Average sensitivity vs. average dose for each schedule is plotted in Fig. 5B. The CT strategy, again, results in a selection for resistant cells over time, but the AT1 strategy, on the other hand, shifts to the lowest dose that still kills the most sensitive phenotypes. After the most sensitive cells are killed, the next most proliferative cells will have a net positive proliferation rate and outgrow due to the low dose. This increased growth then causes a dose increase to keep proliferation in check. So sensitive phenotypes are essentially killed off sequentially with ever increasing doses until the resistant cells take a hold of the invasive edge. At that point, the resistant cells have a small (albeit positive) growth rate with limited chance of control. With the AT2 strategy there is a sharp transition between quick drug-free expansions and MTD-induced contractions. As long as mostly sensitive cells continue to lie on the outer rim, the net decline during treatment is approximately equal to the net increase when the dose is zero, and the tumor can be controlled.

**Figure 5.**
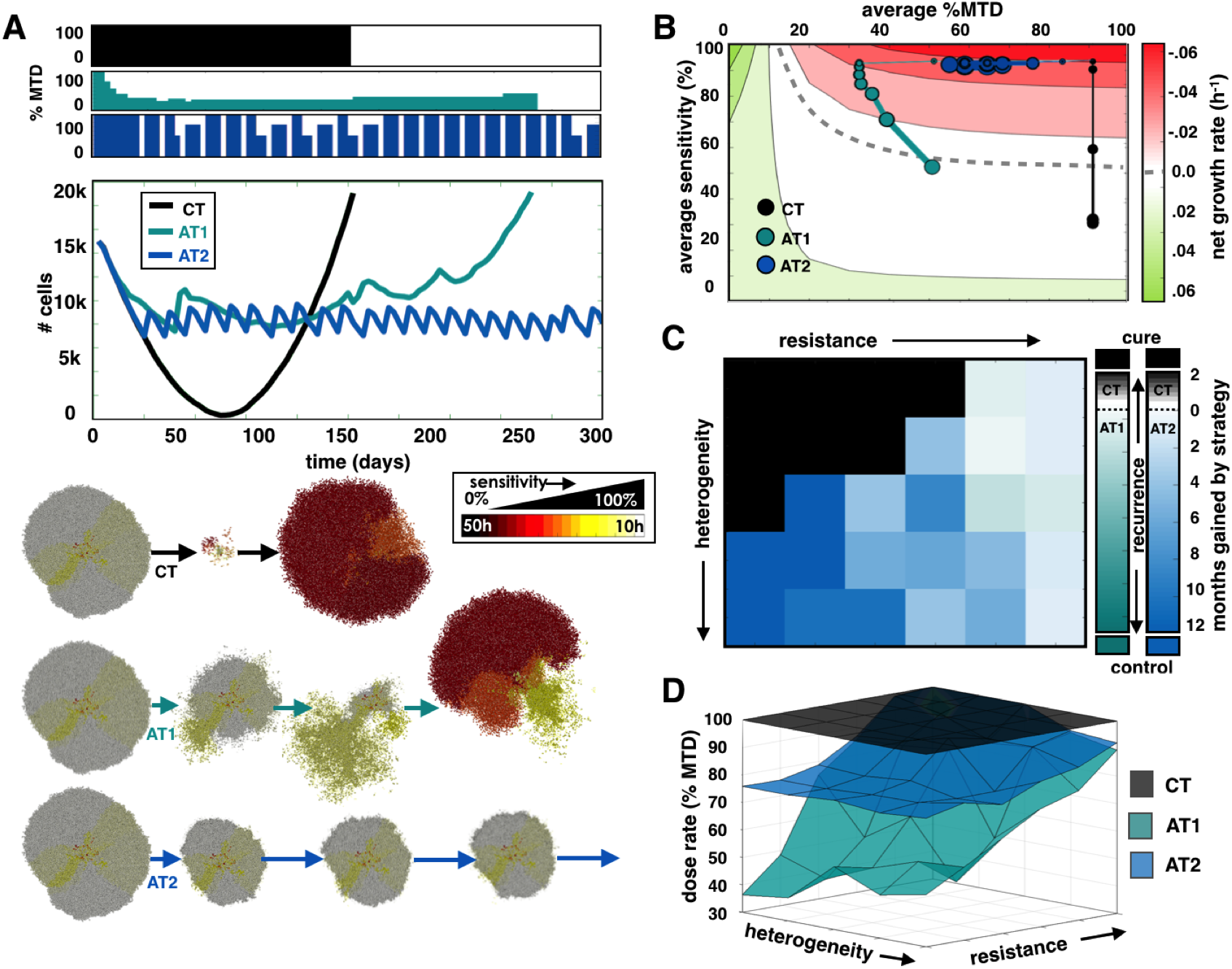
Cell migration (5 μm/h) reduces the efficacy of control by AT. A) Comparing CT, AT1, and AT2 treatments using a tumor from a pre-growth normal distribution of *s*=100% and *σ*_*s*_=25%. Dose schedules, population dynamics, and spatial layout at various time points are shown for each treatment. See link in the Supplement for animations of each of these different treatment strategies. B) Trajectories for average sensitivity vs. average dose over time. C) Winning strategies, and D) Average dose for different initial tumor compositions. See Figure 4 caption for more details on each panel.

A full sweep comparing CT, AT1, and A2 over many tumor compositions is shown in Fig. 5C. We find that while most of the original region cured by CT remains, allowing migration has destroyed the control by AT1 while the regions previously controlled with AT2 are somewhat preserved. The average doses for each strategy are shown in Fig. 5D. The doses are seen to be lower for both AT1 and AT2 in the sensitive region, which was previously controlled (in nonmigrating tumors) by both adaptive strategies. However, control is only achieved with the AT2 strategy. As we increase the migration rate further to 10 μm/h, neither adaptive schedule successfully controls tumors. The time to recurrence using each treatment schedule over the array of tumors is plotted in Fig. S4 for migration rates of 5 and 10 μm/h.

### Treatment vacations help delay recurrence of heterogeneous tumors with phenotypic drift

We have considered treatment responses to tumors that have pre-existing heterogeneity but assume that progeny directly inherit the same phenotypes as the parental cell. However, this ignores the potential impact of subsequent mutations or epigenetic changes that may alter the cell phenotype over generations. Here, we test how phenotypic drift affects the response to treatment strategy, i.e. we allow a cell the opportunity to increase or decrease its proliferation rate upon division (see methods for details).

Starting with a population that is slightly less sensitive but heterogeneous (*s*=80% and *σ*_*s*_=25%), we grow and treat a tumor that can change its proliferation rate (and therefore sensitivity) upon division (Fig. 6A). We find that each treatment strategy eventually fails because now even sensitive cells can become more resistant over generations. However, while CT and AT1 recur at nearly the same time, AT2 is able to control the tumor for around 3 months longer. For the adaptive strategies, resistant regions start to emerge on the tumor's edge, but under the AT2 schedule, the most sensitive cells can regrow during vacation periods and suppress the outgrowth of resistant cells. The cells eventually drift toward more resistant phenotypes, while the sensitive cells are killed, and the mass is separated into individual clumps where the resistant regions exist. We plot the average sensitivity vs. the average dose for each schedule in Fig. 6B. Now, movement along the y-axis occurs not just due to selection but also inheritance of evolving traits. Like all treatment strategies, the drug kills off the most sensitive cells, so the phenotypes eventually end up further into the resistant region, but for CT the recurrent tumor returns almost completely resistant. For AT1, the resistant phenotype emerges faster than the dose modulations can keep up with, but the AT2 schedule, that shifts between extreme doses, better controls both the faster proliferators during treatment and selects against the more resistant cells during vacations. The effect is not sustained for very long.

**Figure 6.**
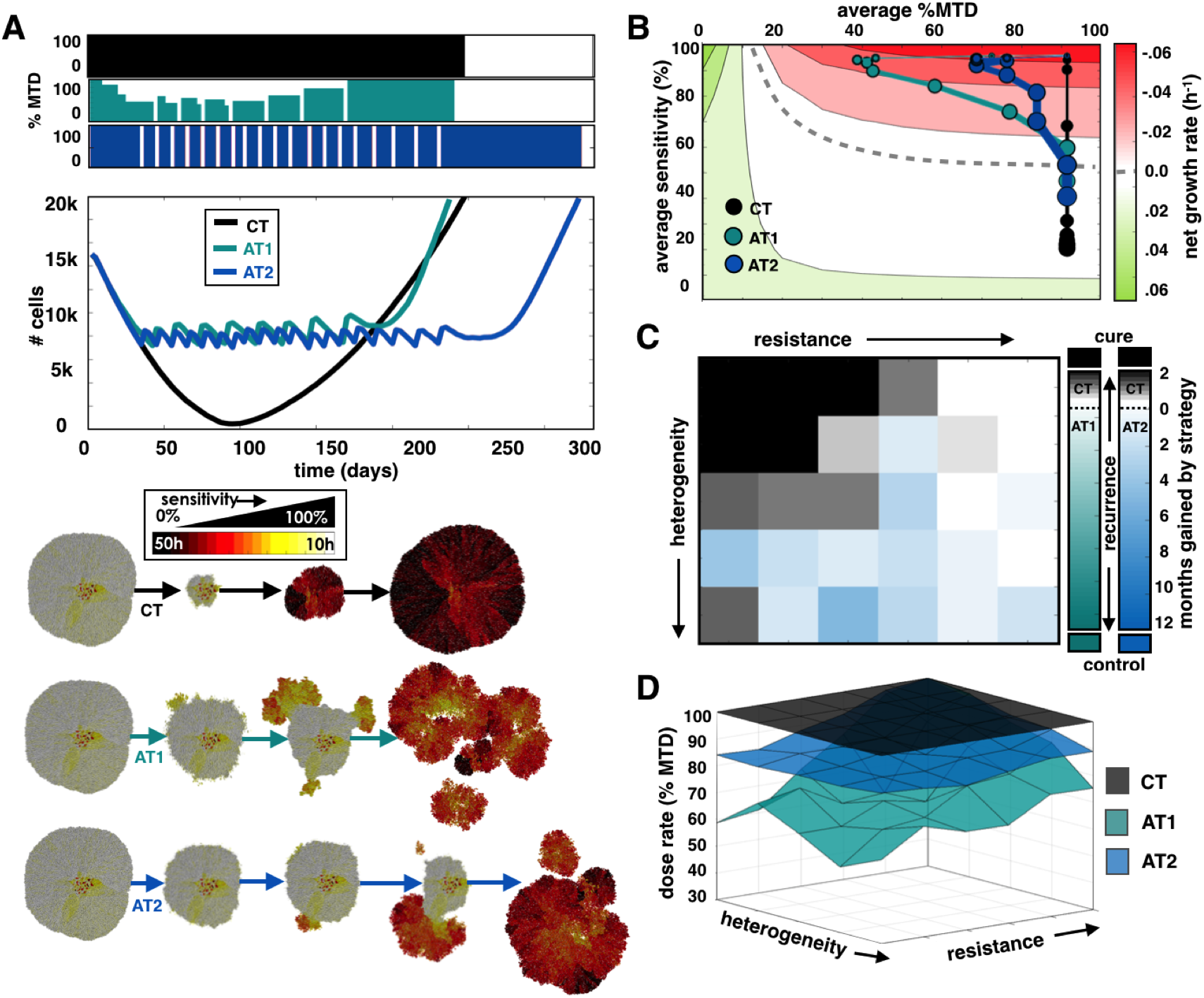
Comparing treatments when tumor cell phenotypes can drift. A) Comparing CT, AT1, and AT2 treatments using a tumor from a pre-growth normal distribution of *s*=80% and *σ*_*s*_=25%. Dose schedules, population dynamics, and spatial layout at various time points are shown for each treatment. See link in the Supplement for animations of each of these different treatment strategies. B) Trajectories for average sensitivity vs. average dose over time. C) Winning strategies, and D) Average dose for different initial tumor compositions. See Figure 4 caption for details on each panel.

The array of different tumor compositions is evaluated to compare CT, AT1, and AT2 (Fig. 6C). The region originally cured by CT is significantly reduced, and the ability to control the tumor through adaptive means is completely eliminated. CT actually does a better job with just a few months gain over AT1 in many regions. But again, the AT2 schedule delays recurrence for longer over the CT schedule for heterogeneous tumor compositions. Regardless, neither treatment sustains the control for long, as the net gain with AT2 is at the most 6 months. The average doses for CT, AT1, and AT2 are shown in Fig. 6D. In the sensitive region, which was controlled when there was no drift, the doses are seen to be higher for both AT1 and AT2. Increasing the drift rate further destroys any control whatsoever using CT, AT1, or AT2. See Fig. S5 for the time to recurrence plots using each treatment schedule over the array of tumors for drift rates of 10% and 100%.

## Discussion

Our simulations demonstrate the necessity of matching any cancer treatment regimen to the corresponding intratumoral evolutionary dynamics. The term “precision medicine” is often applied to strategies that match tumor treatment to specific predictive biomarkers. While this approach increases the probability of success it neglects the reality that virtually all tumor responses are followed by evolution of resistance, and thus, treatment failure. Our results indicate that precision medicine also needs to encompass the complex evolutionary dynamics that govern emergence and proliferation of resistant phenotypes. Here we present the diverse Darwinian interactions that can lead to resistance but also corresponding treatment strategies that can exploit these dynamics to delay the time to progression.

It is clear from our analysis that there is no one-size-fits-all evolutionary strategy. Using a modified off-lattice agent based model, we find conventional MTD application of cancer therapy can cure homogeneous tumors that consist entirely or almost entirely of cells that are sensitive to the applied treatment. While this is improbable at advanced stages, it is observed clinically as some tumors that histologically appear homogeneous, such as testicular cancer and some lymphomas, are frequently cured by conventional chemotherapy. However, decades of clinical experience, consistent with our model predictions, have found cure is not typically achievable in cancers that are highly heterogeneous (e.g. melanoma, lung, breast). Since they already contain cells that are therapy resistant due to genetic, epigenetic or phenotypic properties or environmentally-mediated mechanisms (23).

In tumors that are not curable by standard MTD therapy, our models find that evolution-based strategies that exploit the fitness cost of resistance can delay treatment failure and tumor progression. Recent experimental studies have shown that adaptive therapy can enforce prolonged tumor control (10), however, none of the prior theoretical models have included spatial dynamics and thus assume that the different tumor subpopulations are well mixed (14,15). This is in contrast to radiological and pathological images of tumor that show marked spatial heterogeneity (24–27). Spatial structure can affect the emergence of resistant phenotypes due to space limitation (18,28) and limited drug perfusion (29,30). We present some insight into how spatial context might influence disease control, which reflects similar ideas presented in a recent study with growing bacterial colonies of E. Coli (28). In this study, less fit resistant clones were trapped due to spatial constraints, but high intensity drug exposure led to the competitive release of dormant mutants. Our model shows the same nature of competition, and we are able to test multiple treatment strategies on different tumor phenotypic compositions.

We consider phenotypic heterogeneity over a wide spectrum of tumor compositions. A larger variation in cell phenotypes beyond just sensitive and resistant (with corresponding fitness differences) can drive changes in the dominating population at different doses. By systematically testing an array of different initial phenotypic distributions we can delineate different regions where some schedules work better than others. We also find that some AT strategies are better than others for different situations. For example, with cell migration and phenotypic evolution, we can apply a strategy with less dose modulation and more emphasis on treatment vacations to keep the population sensitive. The extreme dose changes keep the tumor responding quickly during both the growth phase and the treatment phase to either maintain the spatial structure in the case of migration or to prevent the evolution of less sensitive cells in the case of phenotypic drift.

Comparing widely disseminated cancer, where the distribution of tumor cell subpopulations will likely vary among the metastatic sites, to our model, which only considers the response of single tumors, may seem disconnected. It is thus important to ask how much variation in clinical response to CT and AT should be expected. Testing the treatments on a set of tumor metastases with mixed compositions shows that control can still be achieved by using the total numbers of cells across all tumors to represent a systemic measure of burden (Fig. 7). We find that with the CT strategy the more sensitive tumors respond with complete eradication, while the less sensitive tumors eventually recur. In contrast, both AT schedules can still control the disease. While the more sensitive tumors shrink, the less sensitive ones grow in size keeping the total population relatively constant. The full dose schedules are shown in Fig. S6 along with a case where the set of metastases all have the same heterogeneous composition. In the latter situation, the set of tumors respond as if they were single independent tumors. Modeling the spatial distributions of multiple tumors can also be used in conjunction with the periodic systemic measure of tumor burden, which can be tested, calibrated, and validated using blood biomarkers and imaging data. Clearly, there are timescale issues when using data from *in vitro* models (days), *in vivo* models (months), and clinical trials (years) (Fig. 1), but a more rigorous approach to fit data could be done to extract specific growth and response rates for conversion across scales.

**Figure 7.**
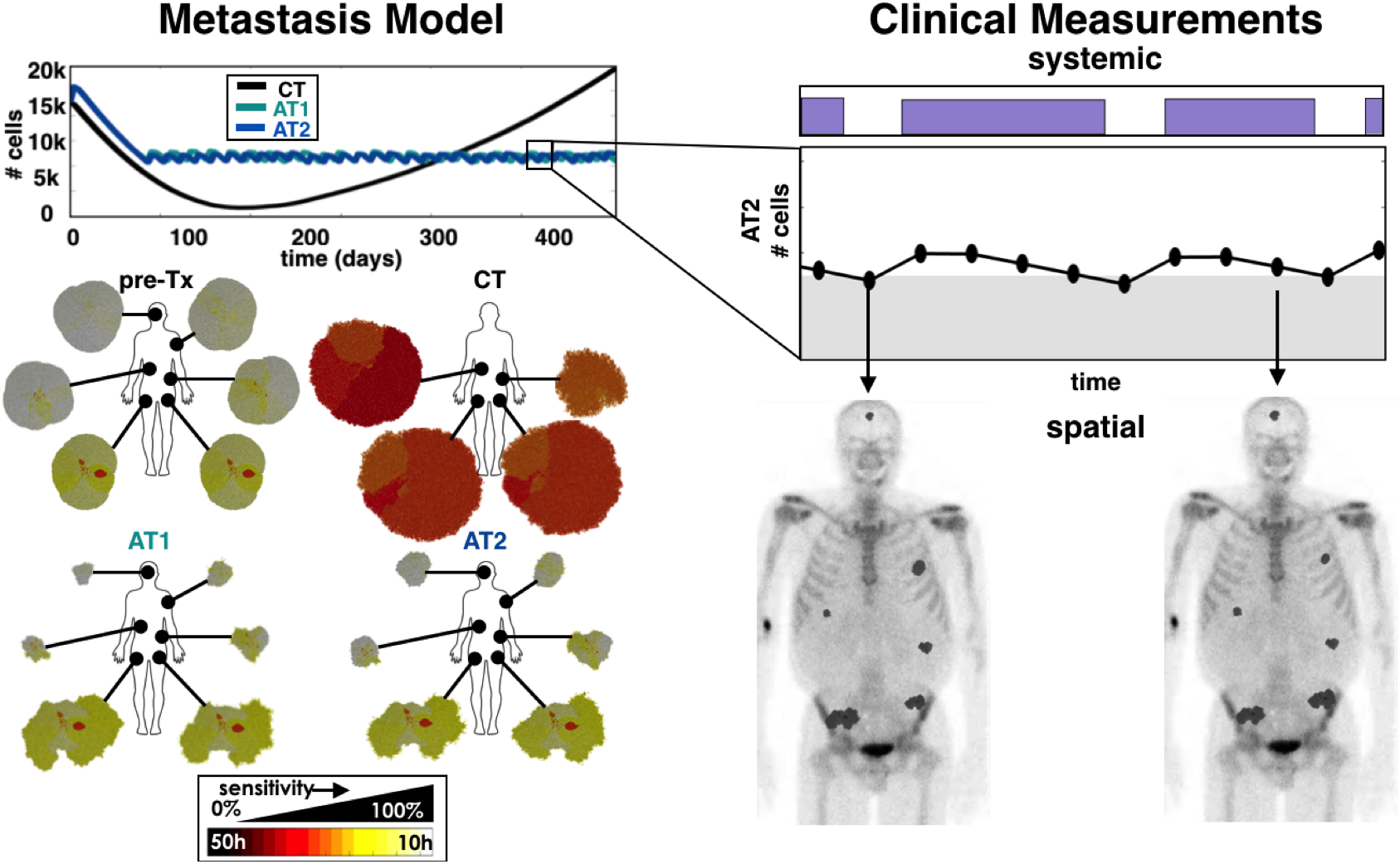
Tumors with dissimilar compositions using CT, AT1 (α=0.25, β=0.05), and AT2 (α=0.50, β=0.10) schedules. The total tumor burden dynamics (the sum of all tumor cells) from all metastatic sites are shown (upper left), and the individual spatial compositions at the end of the simulation are shown (lower left). This model can couple the total systemic tumor burden to the individual spatial distributions found from imaging (right).

The methods for actually measuring tumor burden non-invasively and frequent enough to direct treatment decisions are not well developed for all cancers. In the ongoing adaptive therapy trial on prostate cancer (11), PSA is used as a marker for disease burden. PSA may not be a perfect indicator, but it is readily available, standardized, and utilized (31). Other biomarkers are used for surveillance of various cancers with degrees of specificity and sensitivity (32). As technologies for serum biomarkers, circulating tumor cells, and cell-free DNA continue to be developed (33), we hope this challenge can be addressed to better measure burden in advanced and disseminated cancers through periodic blood draws.

The tumor model presented here is greatly simplified and abstracted to study the impact of spatial intratumoral heterogeneity on tumor progression and how it might be treated using adaptive therapy. We have ignored pharmacokinetics (34), spatially-weighted dose dependence (35,36), microenvironmental influence (37), therapy-induced drug resistance (37), and of course, the 3^rd^ dimension (38). All of these are possible extensions of this work, but we have chosen a simple starting point to understand how a spectrum of cell phenotypes compete for space under different drug strategies. Our work illustrates clearly the importance of using treatment response as a key driver of treatment decisions, rather than fixed strategies. We strongly believe that the future of precision medicine shouldn’t be in the development of new drugs but rather in the smarter evolutionary enlightened application of preexisting ones.

## Acknowledgements

The authors gratefully acknowledge funding from both the Cancer Systems Biology Consortium (CSBC) and the Physical Sciences Oncology Network (PSON) at the National Cancer Institute, through grants U01CA151924 and U54CA193489.

